# A conserved abundant cytoplasmic long noncoding RNA modulates repression by Pumilio proteins in human cells

**DOI:** 10.1101/033423

**Authors:** Ailone Tichon, Noa Gil, Yoav Lubelsky, Tal Havkin Solomon, Doron Lemze, Shalev Itzkovitz, Noam Stern-Ginossar, Igor Ulitsky

**Affiliations:** Department of Biological Regulation, Weizmann Institute of Science, Rehovot 76100, Israel; Department of Molecular Genetics, Weizmann Institute of Science, Rehovot 76100, Israel; Department of Molecular Cell Biology, Weizmann Institute of Science, Rehovot 76100, Israel

## Abstract

Thousands of long noncoding RNA (lncRNA) genes are encoded in the human genome, and hundreds of them are evolutionary conserved, but their functions and modes of action remain largely obscure. Particularly enigmatic lncRNAs are those that are exported to the cytoplasm, including NORAD – an abundant and highly conserved cytoplasmic lncRNA. Most of the sequence of NORAD is comprised of repetitive units that together contain at least 17 functional binding sites for the two Pumilio homologs in mammals. Through binding to PUM1 and PUM2, NORAD modulates the mRNA levels of their targets, which are enriched for genes involved in chromosome segregation during cell division. Our results suggest that some cytoplasmic lncRNAs function by modulating the activities of RNA binding proteins, an activity which positions them at key junctions of cellular signaling pathways.

## Introduction

Genomic studies conducted over the past 15 years have uncovered the intriguing complexity of the transcriptome and the existence of tens of thousands of long noncoding RNA (lncRNA) genes in the human genome, which are processed similarly to mRNAs but appear not to give rise to functional proteins^1^. While some lncRNA genes overlap other genes and may be related to their biology, many do not, and these are referred to as long intervening noncoding RNAs, or lincRNAs. An increasing number of lncRNAs are implicated in a variety of cellular functions, and many are differentially expressed or otherwise altered in various instances of human disease^2^; therefore, there is an increasing need to decipher their modes of action. Mechanistically, most lncRNAs remain poorly characterized, and the few well-studied examples consist of lncRNAs that act in the nucleus to regulate the activity of loci found *in cis* to their sites of transcription^3^. These include the XIST lncRNA, a key component of the X-inactivation pathway, and lncRNAs that are instrumental for imprinting processes, such as AIRN^4^ However, a major portion of lncRNAs are exported to the cytoplasm: indeed, some estimates based on sequencing of RNA from various cellular compartments suggest that most well-expressed lncRNAs are in fact predominantly cytoplasmic^1^.

The functional importance and modes of action of cytoplasmic lncRNAs remain particularly poorly understood. Some lncRNAs that are transcribed from regions overlapping the start codons of protein-coding genes in the antisense orientation can bind to and modulate the translation of those overlapping mRNAs^5^, and others have been proposed to pair with target genes through shared transposable elements found in opposing orientations^6^. Two lncRNAs that are spliced into circular forms were shown to act in the cytoplasm by binding Argonaute proteins (in one case, through ∼70 binding sites for a miR-7 microRNA^7^) and act as sponges that modulate microRNA-mediated repression^7,8^. Such examples are probably rare, as few circRNAs and few lncRNAs contain multiple canonical microRNA binding sites (ref^9^ and IU, unpublished results). It is not clear whether other cytoplasmic lncRNAs can act as decoys for additional RNA-binding proteins through a similar mechanism of offering abundant binding sites for the factors.

The Pumilio family consists of highly conserved proteins that serve as regulators of expression and translation of mRNAs that contain the Pumilio recognition element (PRE) in their 3' UTRs^10^. Pumilio proteins are members of the PUF family of proteins that is conserved from yeast to animals and plants, and whose members repress gene expression either by recruiting 3' deadenylation factors and antagonizing translation induction by the poly(A) binding protein^11^, or by destabilizing the 5' cap-binding complex. The drosophila Pumillio protein is essential for proper embryogenesis, establishment of the posterior anterior gradient in the early embryo, and stem cell maintenance. Related roles were observed in other invertebrates^10^, and additional potential functions were reported in neuronal cells^12^. There are two Pumilio proteins in humans, PUM1 and PUM2^10^, which exhibit 91% similarity in their RNA binding domains, and which were reported to regulate a highly overlapping but not identical set of targets in HeLa cells^13^. Mammalian Pumilio proteins have been suggested to be functionally important in neuronal activity^14^, ERK signaling^15^, germ cell development^16^, and stress response^14^. Therefore, modulation of Pumilio regulation is expected to have a significant impact on a variety of crucial biological processes.

Here, we characterize NORAD – an abundant lncRNA with highly expressed sequence homologs found throughout plancental mammals. We show that NORAD is bound by both PUM1 and PUM2 through at least 17 functional binding sites. By perturbing NORAD levels in osteosarcoma U2OS cells, we show that NORAD modulates the mRNA abundance of Pumilio targets, in particular those involved in mitotic progression. Further, using a luciferase reporter system we show that this modulation depends on the canonical Pumilio binding sites.

## Results

### NORAD is an abundant cytoplasmic lncRNA conserved in mammals

In our studies of mammalian lncRNA conservation we identified a conserved and abundant lincRNA currently annotated as LINC00657 in human and 2900097C17Rik in mouse, and recently denoted as “noncoding RNA activated by DNA damage” or NORAD^17^. NORAD produces a 5.3 Kb transcript that does not overlap other genes (**Figure 1A**), starts from a single strong promoter overlapping a CpG island, terminates with a single major canonical poly(A) site, but unlike most long RNAs is unspliced (**Figure 1B**). Similar transcripts with substantial sequence homology can be seen in EST and RNA-seq data from mouse, rat, rabbit, dog, cow, and elephant. NORAD does not appear to be present in opossum, where a syntenic region can be unambiguously identified based on both flanking genes with no evidence of a transcribed gene in between them, and no homologs could be found in more basal vertebrates. NORAD is ubiquitously expressed across tissues and cell lines in human, mouse, and dog, with comparable levels across most embryonic and adult tissues (**Supplementary Fig. 1)** with the exception of neuronal tissues, where NORAD is more highly expressed. In the presently most comprehensive dataset of gene expression in normal human tissues, compiled by the GTEX project (http://www.gtexportal.org/), the ten tissues with the highest NORAD expression all correspond to different regions of the brain (highest level in the frontal cortex with a reads per kilobase per million reads (RPKM) score of 142), with levels in other tissues varying between an RPKM of 78 (pituitary) to 27 (pancreas). Comparable levels were also observed across ENCODE cell lines, with the highest expression in the neuroblastoma SK-N-SH cells (**Figure 1D**). The high expression levels of NORAD in the germ cells have probably contributed to the large number of closely related NORAD pseudogenes found throughout mammalian genomes. There are four pseudogenes in human that share >90% homology with NORAD over >4 Kb, but they do not appear to be expressed, with the notable exception of the lincRNA transcript HCG11, which is expressed in a variety of tissues but at levels ∼20-times lower than NORAD (based on GTEX and ENCODE data, **Figure 1D**). Due to this difference in expression levels we assume that while most of the experimental methods we used are not able to distinguish between NORAD and HCG11, the described effects likely stem from the NORAD locus and not from HCG11. Using single-molecule *in situ* hybridization (smFISH)^18^ in U2OS cells, we found that NORAD localizes almost exclusively to the cytoplasm (**Figure 1C** and **Supplementary Fig. 2**) and similar cytoplasmic enrichment is observed in other cells lines (**Figure 1D**). The number of NORAD copies expressed in a cell is ∼80 based on the RPKM data and 68±8 based on the smFISH experiments that we have performed on U2OS cells, while 94% of NORAD copies are located in the cytoplasm and 6% are in the nucleus.

**Fig. 1.**
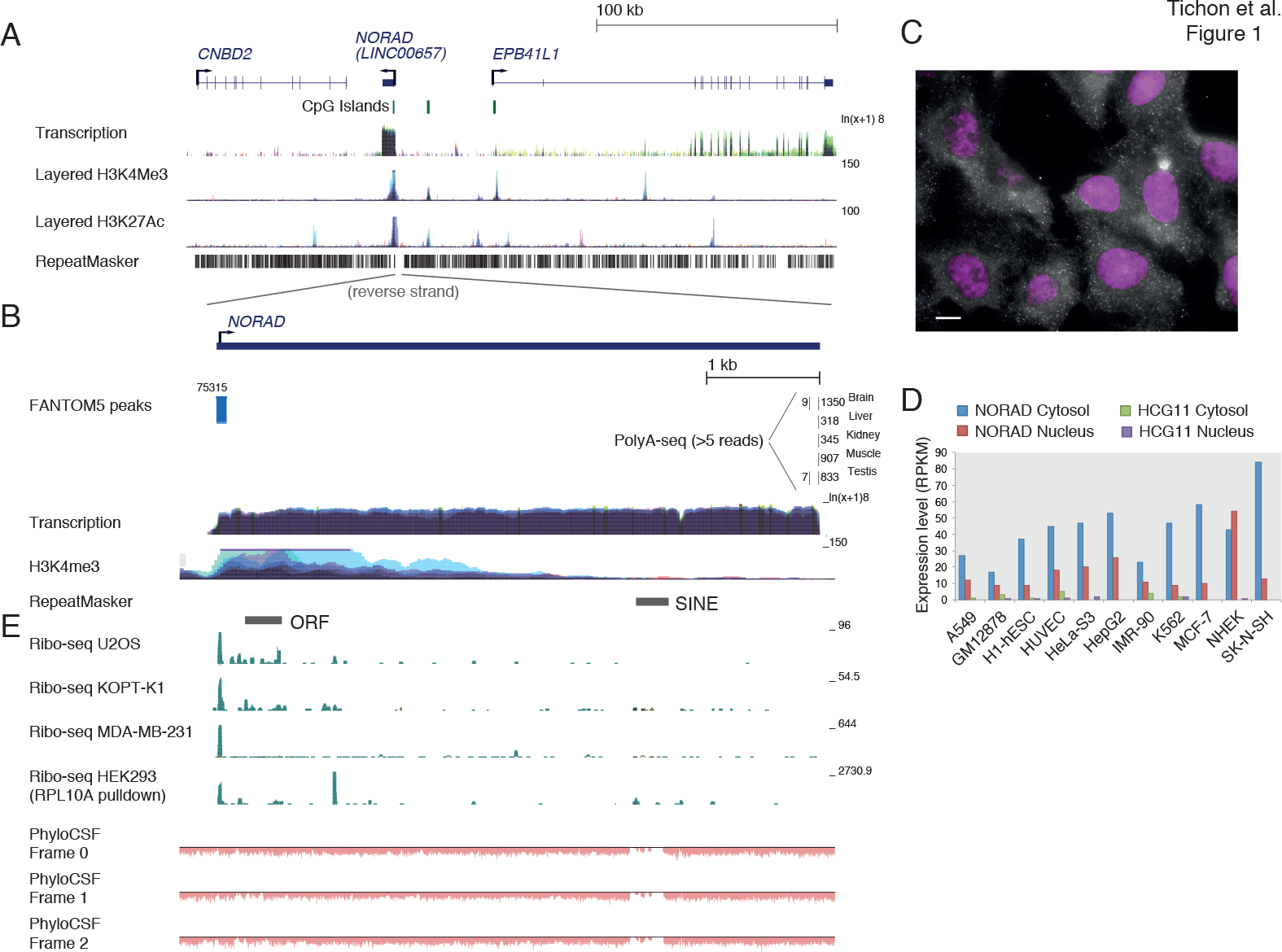
Overview of the human NORAD locus. **(A)** Genomic neighborhood of NORAD. CpG island annotations and genomic data from the ENCODE project taken from the UCSC genome browser. **(B)**Support for the transcription unit of NORAD. Transcription start site information taken from the FANTOM5 project^43^. Polyadenylation sites taken from PolyA-seq dataset^44^. ENCODE datasets and repeat annotations from the UCSC browser. **(C)**Predominantly cytoplasmic localization of NORAD by smFISH. Scale bar 10μm. See Supplementary Fig. 2 for RNA-FISH following NORAD knockdown. **(D)**Estimated copy number of NORAD in 22 independent cells from two independent experiments. **(E)**Expression levels of NORAD and HCG11 in the ENCODE cell lines (taken from the EMBL-EBI Expression Atlas (https://www.ebi.ac.uk/gxa/home). **(E)** Support for the noncoding nature of NORAD. Ribosome protected fragments from various human cell lines (MDA-MB-231^22^, HEK-293^23^, U2OS^24^, KOPT-K1^25^) mapped to the NORAD locus and PhyloCSF^45^ scores. All PhyloCSF scores in the locus are negative.

### NORAD is a bona fide noncoding RNA

NORAD is computationally predicted to be a noncoding RNA by the PhyloCSF (**Figure 1E**) and Pfam/HMMER pipelines, with CPAT^19^ and CPC^20^ giving it borderline scores due to the presence of an open reading frame (ORF) with >100aa (see below) and similarity to hypothetical proteins (encoded by NORAD homologs) in other primates. Therefore, we also examined whether NORAD contains any translated ORFs using Ribo-seq data^21^. When examining ribosome footprinting datasets from diverse human cell lines (MDA-MB-231^22^, HEK-293^23^, U2OS^24^, and KOPT-K1^25^), we did not observe any substantial footprints over any of the ORFs in NORAD, including a poorly conserved 108 aa ORF found close to the 5' end of the human transcript (**Figure 1E**). Interestingly, substantial pileups of ribosome-protected fragments was observed at the very 5' end of NORAD in all Ribo-seq datasets we examined (**Figure 1E** and **Supplementary Fig. 3**), but those did not overlap any ORFs with neither the canonical AUG start codon nor any of the common alternative start codons (**Supplementary Fig. 3**). Additionally, the region overlapping the protected fragments also does not encode any conserved amino acid sequences in any of the frames. We conclude that it is highly unlikely that NORAD is translated into a functional protein under regular growth conditions in those cell types, and the footprints observed in Ribo-seq data result from either a ribosome stalled at the very beginning of a transcript, or from a contaminant footprint of a different ribonucleoprotein (RNP) complex, as such footprints are occasionally present in Ribo-seq experiments^23,26^. It remains possible that NORAD is translated in other conditions and contexts.

### NORAD contains at least 12 structured repeated units

When comparing the NORAD sequence to itself, we noticed a remarkable similarity among some parts of its sequence (**Figure 2A**). Manual comparison of the sequences revealed that the central ∼3.5 kb of NORAD in human, mouse, and other mammalian species can be decomposed into twelve repeating units of ∼300 nt each. Interestingly, these units appear to have resulted from a tandem sequence duplication that occurred at least 100 million years before the split of the eutherian mammals, as when performing pairwise comparisons among repeats from different species, units from different species were more similar to each other than to other units from the same species (data not shown). Overall, the sequences have diverged to a level where there are no sequence stretches that are strictly identical among all the repeats in human. At the core of the most conserved regions within the repeats we identify four sequence and structure motifs (**Figures 2C-E**), some combination of which appears in each of the repeats: (i) one or two PREs (defined by the consensus UGURUAUA); (ii) a short predicted stem-loop structure with four paired bases and a variable loop sequence. The importance of the structure is supported by the preferential A→G and G→A mutations in the second stem-loop that would preserve the stem (**Figures 2D** and **Supplementary Fig. 4**, also detected by EvoFold^27^); (iii) a U-rich stretch of 2-5 bases; (iv) a stem-loop structure with eight or nine predicted base pairs. Further sequence conservation is found upstream and downstream of these motifs. Interestingly, the sequences of some of the repeated units, namely 3-5 and 7-9, appear to be more constrained during mammalian evolution than others (**Figure 2C**), and those units also tend to contain all of the repeat motifs, with more intact sequences and structures (**Figure 2E**).

**Fig. 2.**
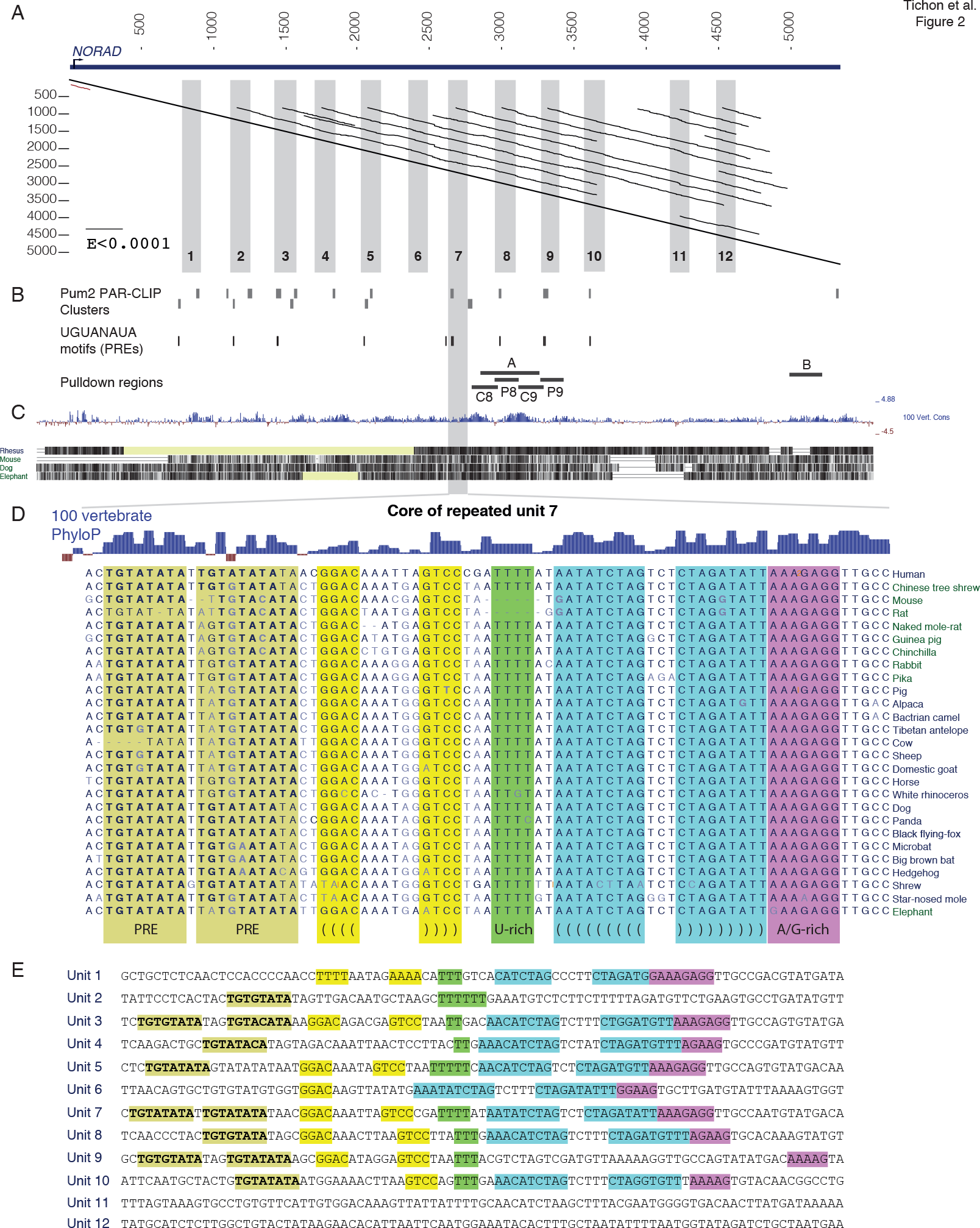
The repeated nature of the NORAD RNA. **(A)** A dotplot computed using plalign^46^ (http://fasta.bioch.virginia.edu/) comparing NORAD with itself. The offdiagonal lines indicate high scoring local alignments between different parts of the sequence. Grey boxes indicate the core of the 12 manually annotated repeated units. **(B)** Clusters identified by PARalyzer^47^ within the NORAD sequence using the PUM2 PAR-CLIP data^28^, positions of UGURUAUA motifs, and regions used for in vitro transcription and pulldown of NORAD fragments. **(C)** Sequence conservation of the NORAD locus, with PhyloP^48^ scores for single-base-level conservation. **(D)** Detailed conservation of the seventh repeated unit. Shaded regions indicate the five motifs present in most repeated units. PREs are Pumilio recognition elements. **(E)** Core sequences of ten of the 12 repeated units, with the same shading as in D.

### NORAD contains multiple functional Pumilio binding sites

In order to identify potential protein binding partners of the repeating units and of other NORAD fragments we first amplified the 8^th^ repeat unit and a region from the 3' end of NORAD, transcribed *in vitro* the sense and antisense of the 8^th^ repeat and the sense of the 3' end region (regions A and B, **Figure 2B**) using the T7 polymerase with biotinylated UTP bases, incubated the labeled RNA with U2OS cell lysate, and subjected the resulting material to mass spectrometry. Among the proteins identified as binding different regions of NORAD (**Supplementary Data 1**) we focus here on two that have predicted binding sites within the repeat units – PUMI and PUM2, the two Pumilio proteins found throughout vertebrates^10^. PUMI and PUM2 proteins were enriched when we performed similar pulldowns followed by Western Blots using the regions within repeats 8 and 9 containing the PREs (Regions P8 and P9 in **Figure 2B**, **Figure 3A**) but not when using adjacent sequences (Regions C8 and C9 in **Figure 2B**, **Figure 3A**) or when a PRE in region P9 has been mutated (**Figure 3B**). To gain additional support for a direct interaction between PUM2 and NORAD, we reanalyzed PAR-CLIP data from HEK-293 cells^28^ and found that PUM2 binds at least 17 sites on NORAD (**Figure 2B**). These experimentally verified sites (all exhibiting T**à**C mutations characteristic of PAR-CLIP and overlapping PREs) overlapped ten of the 11 PREs within repeated units 2-10. It is notable that NORAD has an unusual density of PREs encoded in its sequence – there are 17 non-overlapping instances of the UGURUAUA motifs in NORAD compared to 0.38 expected by chance (P<0.001, see Methods). The number and density of Pumilio motifs within NORAD are higher than those found in all but one human gene (PLCXD1, which has 18 PREs mostly located in transposable elements, compared to 0.12 expected).

**Fig. 3.**
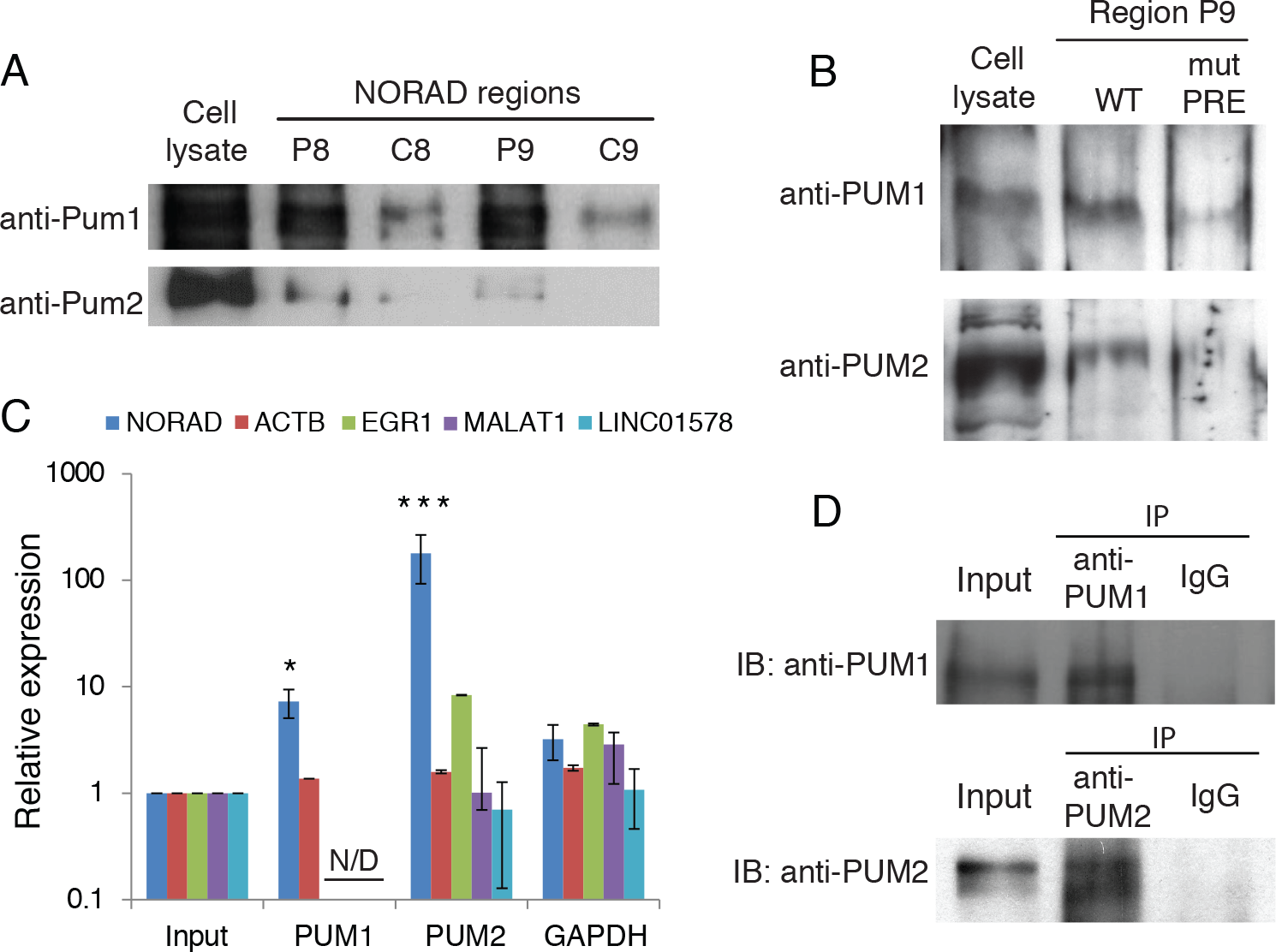
Pumilio proteins bind NORAD. (A) Western blots for PUM1 and PUM2 using the U2OS cell lysate and following pulldowns using the indicated *in vitro* transcribed regions (marked in Figure 2B). (B) Western blots for PUM1 and PUM2 using the U2OS cell lysate and following pulldowns using *in vitro* transcribed RNAs from syntetic oligos with WT or mutated PRE. (C) Recovery of the indicated transcripts in the input and in the indicated IPs. All enrichments are normalized to GAPDH mRNA and to the input sample as described in Materials and Methods. * P<0.05, *** P<0.001. (D) Western blots of the indicated factors in the input and IP samples.

To test whether NORAD also co-precipitates with PUM1 and PUM2 in U2OS cells, we performed RNA Immuno-Precipitation (RIP) of both proteins followed by qRT-PCR, and found a striking enrichment of the NORAD transcript, with a stronger enrichment observed for PUM2 (Methods and **Figure 3C-D**). We conclude that NORAD contains at least 17 confident binding sites for Pumilio proteins, most of which appear in conserved positions within the conserved repeated units. Surprisingly, despite the presence of a large number of bound sites, NORAD levels did not significantly change when PUM1 and PUM2 were over-expressed or knocked-down in U2OS cells (see below), suggesting that it is resistant to substantial degradation by the Pumilio proteins under the tested conditions.

With ∼70 NORAD transcripts per cell (**Figure 1C**) and at least 17 functional PREs (**Figure 2B**), NORAD possesses the capacity to simultaneously bind at least ∼1,200 PUM proteins. Quantitative western blot analysis comparing cell lysates to recombinant proteins expressed in bacteria revealed that PUM1 and PUM2 are expressed at ∼661 and ∼695 copies per cell, respectively (**Supplementary Fig. 5**). The sites offered by NORAD for Pumilio protein binding, as well as the potential interactions made possible by the binding with other NORAD-interacting factors, can be sufficient for eliciting a significant effect on the number of functional Pumilio proteins that are available to act as repressors of their other targets.

### NORAD perturbations preferentially affect Pumilio targets

As PUM1 and PUM2 are reported to affect mRNA stability^11,29^, we next tested if changes in NORAD expression affect the levels of Pumilio targets. We defined Pumilio target genes as those having at least two extra UGUANAUA sites in their 3' UTRs over the number of sites expected given the 3' UTR length of the transcripts. To validate that such genes indeed represent Pumilio targets we knocked down (KD) and over-expressed (OE) PUM1 and PUM2 separately in U2OS cells and observed significant up-regulation of predicted Pumilio targets following KD and down-regulation following OE (**Figure 4A-B**). NORAD was then perturbed using either one of two individual siRNAs (siRNA 1 and siRNA 2, **Supplementary Fig. 6A**) or a pool of four siRNAs (Dharmacon), with the pool yielding ∼4-fold knockdown and individual siRNAs yielding ∼2-fold knockdown (**Supplementary Fig. 6A**). We obtained consistent effects with two independent siRNAs 48 hrs after transfection (**Supplementary Fig. 6B, Supplementary Data 2**), with 51 genes consistently down-regulated by at least 20% after treatment with both siRNAs and 23 genes consistently up-regulated by at least 20%. The stronger knockdown using a pool of siRNAs (**Supplementary Fig. 6A**) resulted in more substantial changes in gene expression – 584 genes were consistently down-regulated by at least 30% in two replicates and 68 genes were consistently up-regulated (**Supplementary Data 2)**. In order to test the consequences of increased NORAD levels, we cloned NORAD into an expression vector where it was driven by a CMV promoter, and transfected the expression vector into U2OS and HeLa cells, which resulted in 2-16 fold up-regulation. Changes following NORAD down-regulation at 24 hr were strongly inversely correlated with the changes observed 24 hr after NORAD over-expression (**Supplementary Fig. 6C** and **Supplementary Data 2**, Spearman *r* =-0.54, P<10^−10^), suggesting that the differential expression was indeed driven by changes in NORAD abundance. Strikingly, Pumilio targets were repressed more than controls when NORAD was downregulated, and their expression levels increased more than controls when NORAD was upregulated in both U2OS and HeLa cells (**Figure 4C-E**). These differences remained significant after controlling for the increased lengths of the 3' UTRs of genes bearing Pumilio motifs (**Supplementary Fig. 7A**) and when considering genes with PUM2 PAR-CLIP clusters in their 3' UTR as determined in HEK-293 cells (these effects were strongest 48 hr after transfection, **Supplementary Fig. 7B**). Genes with multiple PREs were generally more affected than those with fewer sites (**Supplementary Fig. 8A**). Differences between Pumilio targets and controls were observed when considering exon-mapping and not when considering intron-mapping reads, pointing at post-transcriptional regulation^30^ (**Supplementary Fig. 8B**). Lastly, we observed consistent effects in validated PUMI targets^31^ expressed in U2OS cells (**Figure 4F**). These results suggest that hundreds of genes regulated by the two Pumilio proteins are sensitive to NORAD levels, with increased NORAD amounts alleviating repression of Pumilio targets and decreased NORAD amounts increasing repression.

**Fig. 4.**
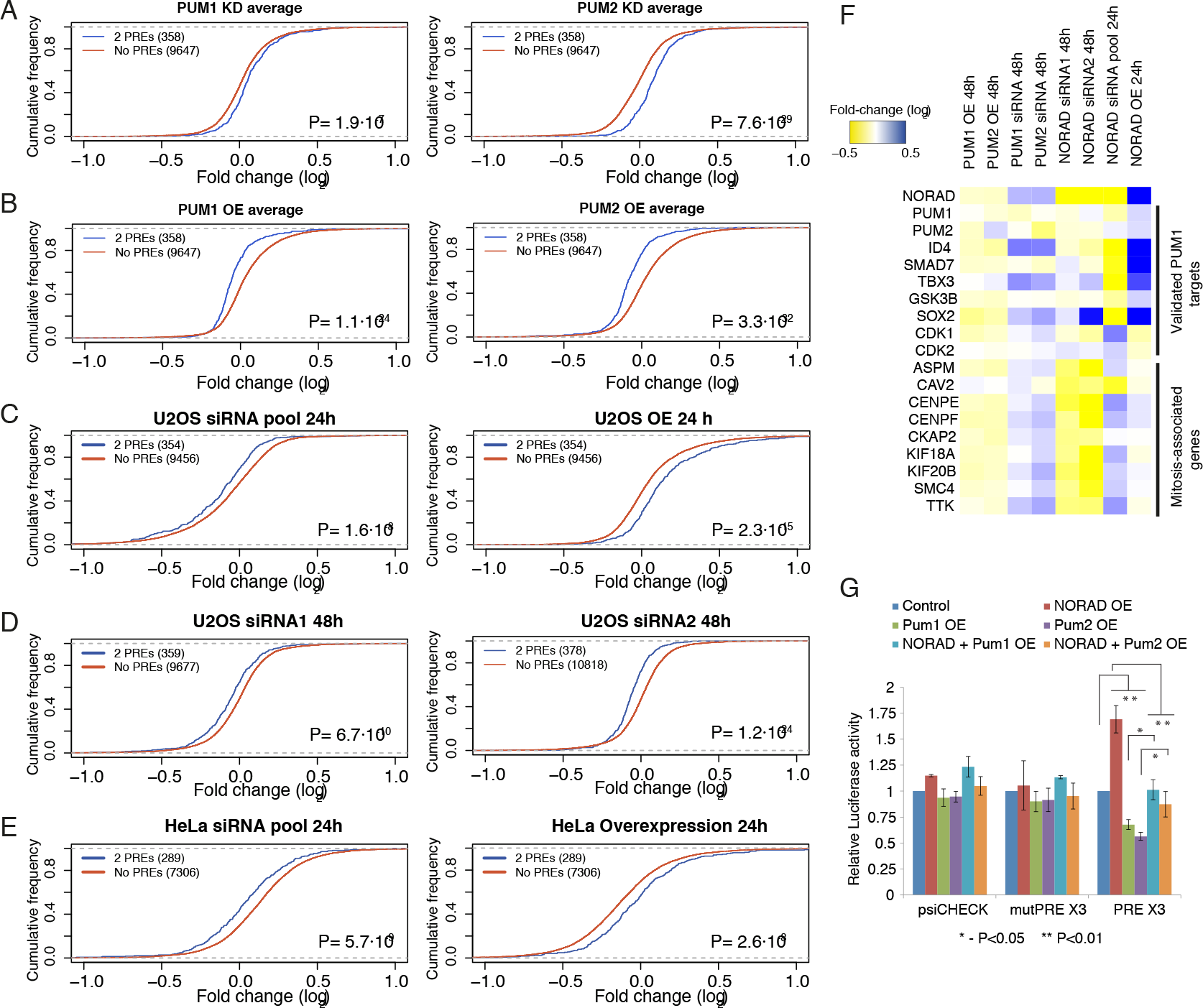
NORAD modulates expression of Pumilio targets. (A-E) Changes in expression of Pumilio targets compared to controls, following the indicated treatment. “2 PREs” are genes that contain at least two canonical PREs over what is expected by chance in their 3' UTRs, “Controls” are those genes that do not contain more sites in their 3' UTRs than expected by chance. (F) Changes in expression of NORAD, PUM1/2, validated targets of PUM1^31^ and genes with annotated roles in the M phase of the cell cycle and/or the mitotic spindle following the indicated perturbations. (G) Changes in luciferase activity measured from the indicated vectors following overexpression of the indicated genes and combinations.

When we inspected the Gene Ontology annotations enriched in the different sets of genes responsive to NORAD perturbations, after correction for multiple testing using TANGO^32^, the only significantly enriched group were genes bound by PUM2 in the PAR-CLIP data and down-regulated 48 hr after NORAD knockdown. These genes were enriched with categories associated with cell cycle and mitosis, including “M phase of the cell cycle” (8 genes; P= 6.4 × 10^−6^) and “Spindle” (8 genes; *P* = 1.2 × 10^−7^). Interestingly, these genes were not substantially affected at 24 hr after NORAD knockdown or over-expression (**Figure 4F**), and enrichments of NORAD targets were also significant when compared to all PUM2-bound targets, suggesting a cumulative, and perhaps cell-cycle-dependent, effect of NORAD perturbation on Pumilio targeting of genes involved in mitosis. These results are consistent with the chromosomal instability and mitotic defects observed in other cell types following TALEN-mediated deletion of NORAD^17^.

As Pumilio proteins may affect translation in addition to their effects on mRNA stability, we evaluated the translational consequences of NORAD perturbation after 48 hr using Ribo-seq^33^. Consistent with the RNA-seq data, the number of translating ribosomes on Pumilio targets was reduced following NORAD KD (**Supplementary Fig. 9A**). However, when normalizing for changes in mRNA levels, translation efficiency of Pumilio targets did not appear to be preferentially affected (**Supplementary Fig. 9B**), suggesting that the main effects of NORAD on Pumilio targets are through effects on mRNA stability rather than translation. This observation is consistent with reports that the mechanism of action of Pumilio proteins is through interaction with deadenylation complexes^11,29^ that can first affect protein translation, but eventually results in mRNA decay.

### NORAD regulation is dependent on the canonical PREs

In order to test whether regulation of Pumilio targets depends on canonical PREs, we utilized a luciferase reporter vector containing three strong PREs as well as a control reporter with mutated sites, in which the three UGUACAUA motifs were mutated to ACAACATA (mutPRE)^11,29^. As expected, over-expression of PUM1 or PUM2 proteins in U2OS cells led to increased repression in a PRE-dependent manner (**Figure 4G**). Over-expression of NORAD, on the other hand, alleviated the repression of the PRE-containing luciferase mRNA, without affecting luciferase containing mutPRE elements. Simultaneous over-expression of NORAD and the Pumilio proteins abrogated both effects, consistent with an effect of NORAD on the ability of Pumilio proteins to repress their targets (**Figure 4G**). Knockdowns of NORAD or PUM1/2 failed to yield a consistent effect on luciferase activity (**Supplementary Fig. 10A**), possibly because of the limited knockdown efficiency using siRNAs (**Supplementary Fig. 10B**) or through feedback regulation of PUM1/2 on their own mRNA (see Discussion). Overall, these results indicate that the NORAD-dependent changes in abundance of Pumilio targets are likely mediated through canonical PREs.

## Discussion

To our knowledge, NORAD comprises the first example of a lncRNA that contains multiple highly conserved consensus binding sites for an RNA binding protein (RBP), and that is required for proper regulation of the RBP targets at physiological levels. One particularly interesting question that remains open is the functional importance and roles of the other conserved elements found in the NORAD repeats, and in particular the two predicted hairpin structures, as such conserved secondary structures are generally rarely detectable in lncRNAs^1^. It is possible that these structural elements serve as binding sites for other RBPs, whose binding may either facilitate the binding of PUM1 and PUM2 to NORAD or affect PUM1/2 protein stability or activity. We note that while the overall number of binding sites offered by NORAD for PUM1 and PUM2 (∼1200) is comparable to the Pumilio abundance in U2OS cells, which we estimate at ∼650 copies per cell for each of PUM1 and PUM2 (**Supplementary Fig. 5**), these sites are outnumbered by the sites present in other expressed mRNAs, and therefore it is possible that NORAD does not merely titrate Pumilio proteins away from their other targets but rather induces a changes in their activity, potentially by serving as a scaffold for interaction of Pumilio proteins with other factors. Potentially interesting candidates for interacting with NORAD repeats that were identified in the mass spectrometry analysis are known RBPs such as IGF2BP1/2/3, XRN2, and PABPN1. In addition we observed that the interferon response pathway proteins IFIT1/2/3/5 and their downstream companion PKR could bind NORAD sequence. IFIT proteins were observed to bind the antisense of the NORAD 8^th^ repeat unit, suggesting that they may recognize a structural element rather than a primary sequence within the repeat, whereas PUM1 and PUM2 bound only the sense sequence, consistent with its known sequence specificity. We were so far unable to substantiate interactions with IFIT1 and PKR by reciprocal pulldown experiments, but if this interaction is indeed specific it would link NORAD to the reported functions of Pumilio in viral response – PUM1 and PUM2 were shown to be functionally stimulated after migration into Stress Granules upon viral infection^34^ – an event that induces the interferon pathway.

While this manuscript was under review, Mendell and colleagues described a role for NORAD and PUM2 in ensuring chromosomal segregation fidelity in various human cells. Further studies will be required in order to uncover the full spectrum of physiological consequences of the regulation of Pumilio targets by NORAD, but the enrichment of cytokinesis-related genes among the Pumilio targets that are sensitive to NORAD levels suggests that NORAD may modulate regulation of chromosomal segregation during mitosis by Pumilio, and might even affect the conserved roles of Pumilio in regulating asymmetric cell divisions during embryonic development. An intriguing question is whether the relatively high levels of NORAD in U2OS cells correspond to a basal state, in which NORAD exerts a minimal effect on PUM1/2 that is increased when stimuli increase NORAD expression, or to a state where NORAD actively buffers substantial regulation by PUM1/2. Most results point to the former scenario, as relatively modest over-expression of NORAD resulted in stronger effects on Pumilio activity than its knockdown. Another possibility suggested by the enrichment of cell-cycle regulated genes among the most prominent NORAD/Pumilio targets is that this regulation is cell-cycle dependent.

Another interesting question is whether NORAD affects Pumilio target regulation through binding a substantial number of functional PUM proteins through its numerous binding sites or by transient binding that alters PUM stability or activity. Answering this question is complicated by the negative autoregulation of PUM2, which is binding their own 3' UTRs^28^. We did not observe consistent and strong effects of NORAD perturbations on PUM1/2 mRNA or protein level but it is possible that those effects are masked by the feedback regulation. If, for instance, NORAD binding facilitates Pumilio protein degradation, we are expecting increased PUM1/2 production that may result in unaltered PUM1/2 protein levels but reduced availability of functional Pumilio proteins in the cells.

## Materials and methods

### Cell culture

Human cell lines U2OS (osteosarcoma) and HeLa (cervical carcinoma) were routinely cultured in DMEM containing 10% fetal bovine serum and 100 U penicillin/0.1 mg/ml streptomycin at 37 °C in a humidified incubator with 5% CO_2_.

### Plasmids and siRNAs

Plasmid transfections were performed using PolyEthylene Imine (PEI)^35^ (PEI linear, Mr 25000 from Polyscience Inc). In order to overexpress NORAD, the full transcript of the lincRNA was amplified from human genomic DNA (ATCC NCI-BL2126) using the primers TGCCAGCGCAGAGAACTGCC (Fw) and GGCACTCGGGAGTGTCAGGTTC (Rev), and cloned into a ZeroBlunt TOPO vector (Invitrogen), and then subcloned into the pcDNA3.1(+) vector (Invitrogen). PUM1 and PUM2 were over-expressed using pEF-BOS vectors^34^ (a kind gift of Prof. Takashi Fujita). As controls in over-expression experiments we used pBluescript II KS+ (Stratagene). Plasmids were used in the amount of 0.1 μg per 100,000 cells in 24 well plates for 24 h before cells were harvested. The luciferase experiments employed the following plasmids: pGL4.13; psiCheck-1 containing 3X wild type PRE, which is underlined in the following sequence, 5’-TTGTTGTCGAAAAT<u>TGTACATA</u>AGCCAA; psiCheck-1 containing 3X mutated PREs: 5'-TTGTTGTCGAAAAT**<u>ACA</u>**<u>ACATA</u>AGCCAA and psiCheck-1 with no PRE, all previously described^11,29^ (a kind gift of Dr. Aaron Goldstrohm). pGL4.13 was used in the amount of 5ng per 20,000 cells in 96 well plates while the different psiCheck-1 plasmids were used in the amount of 15ng per 20,000 cells in 96 well plates. Transfection time was 48h prior to further experimental procedures.

Gene knockdown was achieved using siRNAs directed against NORAD, PUM1, and PUM2 genes (all from Dharmacon, **Supplementary Table 1**), while as control we used the mammalian non-targeting siRNA (Lincode Non-targeting Pool, Dharmacon), at final concentration of 50nM for 24h or 48h prior to further experimental procedures. The transfections into U2OS cells were conducted using the PolyEthylene Imine.

siRNA transfection into HeLa cells were conducted using 100 nM siRNA and Dharmafect (Dharmacon) transfection reagent and using siRNA buffer only as a control, and transfection of pCDNA3.1-NORAD was into HeLa cells was peformed using Lipofectamine 2000.

### Real-time PCR analysis of gene expression

Total RNA was isolated using TRI reagent (MRC), followed by reverse transcription using an equal mix of oligo dT and random primers (Quanta), according to the manufacturer’s instructions. For determination of all genes levels real-time PCR was conducted using Fast SYBR qPCR mix (Life technologies). The primer sets used for the different genes are listed in **Supplementary Table 2**. The assays contained 10-50 ng sample cDNA in a final volume of 10 μl and were run on AB qRT-PCR system ViiA 7 (Applied Biosystems). All genes expression levels in the different treatments are represented relative to their relevant control (ΔCt) and normalized to GAPDH gene levels (ΔΔCt).

### Fluorescent In-Situ Hybridization

Probe libraries were designed according to Stellaris guidelines and synthetized by Stellaris as described in Raj et al^18^. Libraries consisted of 48 probes 20 nt each, complementary to the NORAD sequence according to the Stellaris guidelines (**Supplementary Table 3**). Hybridizations were done overnight at 30°C with Cy5 labeled probes at a final concentration of 0.1ng/μl. DAPI dye (Inno-TRAIN Diagnostik Gmbh) for nuclear staining was added during the washes. Images were taken with a Nikon Ti-E inverted fluorescence microscope equipped with a 100 × oil-immersion objective and a Photometrics Pixis 1024 CCD camera using MetaMorph software (Molecular Devices, Downington, PA). The image-plane pixel dimension was 0.13 μm. Quantification was done on stacks of 4-12 optical sections with Z-spacing of 0.3 μm. Dots were automatically detected using a custom Matlab program, implementing algorithms described in Raj et al^18^. Briefly, the dot stack images were first filtered with a three-dimensional Laplacian of Gaussian filter of size 15 pixels and standard deviation of 1.5 pixels. The number of connected components in binary thresholded images was then recorded for a uniform range of intensity thresholds and the threshold for which the number of components was least sensitive to threshold selection was used for dot detection. Automatic threshold selection was manually verified and corrected for errors. Background dots were detected according to size and by automatically identifying dots that appear in more than one channel (typically <1% of dots) and were removed.

### Biotin pulldown assay

Templates for in-vitro transcription were generated by amplifying the desired sequences from cDNA or from synthetic oligos, adding the T7 promoter to the 5’ end for sense and 3’ end fort he antisense sequence (See Supplementary Table 2 for primer sequences). In addition, protein pulldown was performed using an oligo with the sequence of repeat #9 (5‘-GTCTGCATTTTCATTTACTGTGCTGTGTATATAGTGTATATAAGCGGACATAGGAGTCCTAATTTACGTCTAGTCGATGTTAAAAAGGTTGCCAGTATATGACAAAAGTAGAATTAGTAAACTACTACATTGAGTACACTTTGTGTTAAAATTCATAGGGA) and an oligo that contains a mutation in its PRE (5‘- GTCTGCATTTTCATTTACTGTGCT**<u>ACA</u>**TATATAGTGTATATAAGCGGACAT AGGAGTCCTAATTTACGTCTAGTCGATGTTAAAAAGGTTGCCAGTATATGACAAAAGTAGAATTAGTAAACTACTACATTGAGTACACTTTGTGTTAAAATTCATAGGGA) using the primers from Supplementary Table 2. Biotinylated transcripts were produced using the MEGAscript T7 in-vitro transcription reaction kit (Ambion) and Biotin RNA labeling mix (Roche). Template DNA was removed by treatment with Dnasel (Quanta). Protein extract was prepared by lysing the cells in buffer containing 20mM Tris-HCl at pH 7.5, 150 mM NACl, 1.5 mM MgCl_2_, 2mM DTT, 0.5% Na-Deoxycholate, 0.5% NP-40) incubating on ice for 15 min followed by centrifugation at 15000 RPM at 4°C for 20 min. Then, 0.5-2mg of protein were pulled-down by incubation with the 2-20pmole biotynylated transcripts. The pulldown products were analyzed by mass spectrometry and western blots. For the mass spectometry the formed RNA-protein complexes were precipitated by Streptavidin-sepharose high performance beads (GE Healthcare), and then proteins were then resolved on a 4-12% Express Page gradient gel (GeneScript) and visualized by silver staining. The entire lane was extracted and analyzed using Mass spectrometry analysis as previously described^36^. For the western blot assays, formed RNA-protein complexes were precipitated by Streptavidin magnetic beads (Invitrogen), seperated on a 10% SDS-PAGE gel, and incubrated with anti-PUM1 or anti-PUM2 antibodies (Bethyl).

### RNA Immunoprecipitation (RIP)

Immunoprecipitation (IP) of endogenous RNP complexes from whole-cell extracts was performed as described by Yoon et al^37^. In brief, cells were incubated with lysis buffer (20 mM Tris-HCl at pH 7.5, 150 mM NACl, 1.5 mM MgCl_2_, 2mM DTT, 0.5% Na-Deoxycholate, 0.5% NP-40, complete protease inhibitor cocktail (Sigma) and 100unit/ml RNase inhibitor (EURx)) for 15 min on ice and centrifuged at 13,000 RPM for 15 min at 4°C. Part of the supernatants was collected for the total cell lysate input. The rest, containing 1-2 mg protein extract, was incubated for 2-3 hours at 4°C in gentle rotation with protein A/G magnetic beads (GeneScript) that were pre-washed and coated with antibodies against GAPDH (SantaCruz Biotechnology), PUM1 and PUM2 (Bethyl) at 4°C in gentle rotation over-night. As a negative control, we incubated the magnetic beads-antibodies complexes with lysis buffer. Afterwards, the beads were washed five times with the lysis buffer, each time separated by magnetic force. The remaining mixture of magnetic beads-antibodies-protein-RNA complexes were separated as half were mixed with sample buffer and boiled at 95C for 5 minutes for further analysis by Western blot and the other half was incubated with 1mg/ml Proteinase K for 30min at 37C with gentle shaking to remove proteins. The remaining RNA was separated by Trizol. The RNPs isolated from the IP materials was further assessed by RT-qPCR analysis as follows: 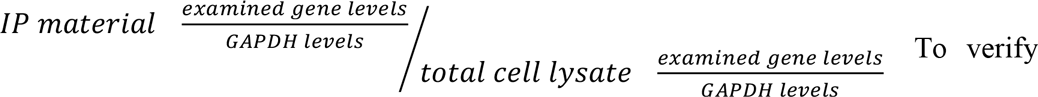that the desired protein was indeed immune-precipitated the Western blot assay was employed as described above.

### Ribosome profiling

48h after transfection of the U2OS with siRNAs, cylcoheximide treatments were carried out as previously described^21^. Cells were lysed in lysis buffer (20mM Tris 7.5, 150mM KCl, 5mM MgCl2, 1mM dithiothreitol, 8% glycerol) supplemented with 0.5% triton, 30 U/ml Turbo DNase (Ambion), and 100μg/ml cycloheximide, and ribosome-protected fragments were then generated, cloned, and sequenced as previously described^21^.

### RNA-seq and data analysis

Strand-specific mRNA-seq libraries were prepared from U2OS cells using the TruSeq Stranded mRNA Library Prep Kit (Illumina) and sequenced on a NextSeq 500 machine to obtain at least 23 million 75 nt reads. Strand-specific mRNA-seq libraries for HeLa cells were prepared as described^38^. All sequencing data has been deposited to the GEO database (Accession GSEXXXX). Reads were aligned to the human genome (hg19 assembly) using STAR Aligner^39^, and read counts for individual genes (defined as overlapping sets of RefSeq transcripts annotated with their Entrez Gene identifier) were counted using htseq-count^40^ and normalized to reads per million aligned reads (RPF). For counting intron-mapping reads, htseq-count was used to count reads mapping to the whole gene locus, and the exon-mapping reads were then subtracted for each gene. Only genes with an average RPM of at least 50 normalized reads across the experimental conditions were considered, and fold changes were computed after addition of a pseudo-count of 0.1 to the RPM in each condition. The raw read counts and the computed fold-changes appear in Supplementary Data 2.

### Sequence analyses

Whole genome alignments were obtained from the UCSC genome browser. Expected numbers of PREs were computed by applying dinucleotide-preserving permutations to the sequences and counting motif occurrences in the shuffled sequences. 3’ UTR-length-matched control targets were selected by dividing the genes into ten bins based on 3' UTR lengths and randomly sampling the same numbers of genes not enriched with Pumilio target sites as the number of genes enriched with sites from each bin.

### Luciferase assays

The activity of Pumilio was determined by Luciferase assay as previously described^41^. Briefly, 20,000 Cells were plated in a 96-well plate. After 24 hr cells were cotransfected with pGL4.13 as an internal control and with the indicated psiCheck plasmids. In addition, the cells were transfected with the various siRNAs or plasmids (as described above). After 48 hr, luciferase activity was recorded using the Dual-Glo Luciferase Assay System (Promega) in the Micro plate Luminometer device (Veritas). A relative response ratio (RRR), from RnLuc signal/FFLuc signal, was calculated for each sample. Percent of change is relative to the control siRNA or control plasmid.

### Determination of copy number of PUM1 and PUM2 in U2OS cells

PUM1 and PUM2 were expressed in bacteria. Briefly, PUM1 and PUM2 cDNA were cloned into a modified version of pMal-C2 expression vector (a kind gift from the laboratory of Prof. Deborah Fass) by restriction free cloning resulting in a MBP-6His- PUM constructs. The plasmids were transformed into Rosetta-R3 bacteria (Novagen). Bacteria were grown in 15ml 2YT media in the presence of 100μg/ml Ampicillin and 50μg/ml Chloramphenicol to OD600≈0.6. Recombinant protein expression was induced for 18hrs at 16°c by 500μM IPTG.

Bacterial pellet was resuspended in 5ml of lysis buffer B (100mM NaH_2_PO_4_, 10mM Tris, 8M Urea, pH8) and incubated on a rotating shaker for 90 minutes at room temperature. The extract was cleared by centrifugation (10000xg, 20°c for 30min). Cleared extract was incubated for 60min with 1ml of 50% Nickel beads slurry (NiNTA His**·**Bind Rasin, Novagen) and the extract bead mix was loaded onto an empty column. The column was washed twice with wash buffer C (100mM NaH_2_PO_4_, 10mM Tris, 8M Urea, pH6.3) and the bound proteins were eluted 4 times in 500μl of elution buffer D (100mM NaH_2_PO_4_, 10mM Tris, 8M Urea, pH5.9) followed by 4 times in 500μl of elution buffer E (100mM NaH_2_PO_4_, 10mM Tris, 8M Urea, pH4.5). Sample of each fraction was run on SDS-PAGE and analyzed by Coomasie blue staining. To determine the quantity of PUM1 and PUM2 copies per cell we calibrated a standard curve using the purified bacterial expressed PUM proteins and then plotted the protein expression levels from a lysate extracted from a measured number of cells.

### Statistics

All results are represented as an average ± SEM of at least three independent experiments. Statistics was performed as Student’s t-test, Wilcoxon rank-sum test or Anova with Tuckey’s post hoc test for three or more groups to be compared. In all results * p<0.05, ** p<0.01, *** p<0.001. Plots were prepared using custom R scripts. Gene Ontology enrichment analysis was performed using the WebGestalt server^42^ and corrected for multiple testing using TANGO^32^, using all the expressed genes as background set and Benjamini-Hochberg correction for multiple testing.

## Acknowledgements

We thank members of the Ulitsky lab for useful discussions and comments on the manuscript. I.U. is incumbent of the Sygnet Career Development Chair for Bioinformatics and recipient of an Alon Fellowship. Work in the Ulitsky lab is supported by grants to I.U. from the European Research Council (Project “lincSAFARI”), Israeli Science Foundation (1242/14 and 1984/14), the I-CORE Program of the Planning and Budgeting Committee and The Israel Science Foundation (grant no 1796/12), the Minerva Foundation, the Fritz-Thyssen Foundation, and by a research grant from The Abramson Family Center for Young Scientists.

